# Decoding working memory information from persistent and activity-silent neurons in the primate prefrontal cortex

**DOI:** 10.1101/2023.07.25.550371

**Authors:** Lilianna Thrower, Wenhao Dang, Rye G. Jaffe, Jasmine D. Sun, Christos Constantinidis

## Abstract

Persistent activity of neurons in the prefrontal cortex has been thought to represent the information maintained in working memory, though alternative models have recently challenged this idea. Activity-silent theories posit that stimulus information may be maintained by the activity pattern of neurons that do not produce firing rate significantly elevated about their baseline during the delay period of working memory tasks. We thus tested the ability of neurons that do and do not generate persistent activity in the prefrontal cortex of monkeys to represent spatial and object information in working memory. Neurons that generated persistent activity represented more information about the stimuli in both spatial and object working memory tasks. The amount of information that could be decoded from neural activity depended on the choice of decoder and parameters used but neurons with persistent activity outperformed non-persistent neurons consistently. Although averaged across all neurons and stimuli, firing rate did not appear clearly elevated above baseline during the maintenance of neural activity particularly for object working memory, this grant average masked neurons that generated persistent activity selective for their preferred stimuli, which carried the majority of information about the stimulus identity. These results reveal that prefrontal neurons with generate persistent activity constitute the primary mechanism of working memory maintenance in the cortex.

**NEW AND NOTEWORTHY:** Competing theories suggest that neurons that generate persistent activity or do not are primarily responsible for the maintenance of information, particularly regarding object working memory. While the two models have been debated on theoretical terms, direct comparison of empirical results have been lacking. Analysis of neural activity in a large database of prefrontal recordings revealed that neurons that generate persistent activity were primarily responsible for the maintenance of both spatial and object working memory.

## INTRODUCTION

The ability to maintain and manipulate information in memory over a period of seconds is commonly referred to as working memory (1). Early neurophysiological experiments in non-human primates identified neurons that not only respond to sensory stimuli but remain active during a period after the stimuli were no longer present, thus providing a neural correlate of working memory (2). This persistent activity has since been widely replicated, including in human recordings (3). Computational models typically simulate persistent activity through neural networks with recurrent connections between units with similar tuning for spatial location (4). Activation in the network behaves as a continuous attractor (5), with a stimulus generating a bump (peak) of activity in the network. This activity bump is maintained through the delay period, but may drift randomly, with the peak at the end of the delay period determining the location that is ultimately recalled by the subject, hence the term, bump attractor (6). Drifts in neuronal activity thus account for deviations of behavior.

Persistent activity is not universally accepted as the neural correlate of working memory (7–9). Alternative mechanisms have been proposed that do not depend on persistent activity to represent information during the delay period of working memory tasks, including “activity-silent” models, which suggest that information can be decoded from the PFC without generation of elevated persistent activity across the population For example, experimental studies have shown that the identity of a stimulus maintained in memory can be decoded from a population of neurons whose activity is not elevated above baseline levels of firing rate, but whose pattern of activity may vary dynamically over time, with different neurons becoming active at different time points Computational models have also been proposed that could account for such changes via synaptic strength modification instead of spike generation (12, 13). Moreover, Recurrent Neural Networks have been used successfully to model complex cognitive tasks, including those related to working memory (14–17), demonstrating that it is possible to perform working memory tasks by virtue of rapid changes in synaptic weights after a stimulus presentation.

We were therefore motivated to compare the predictions of the bump-attractor and activity-silent models, using a database of recordings obtained from the prefrontal cortex of monkeys as they performed spatial and object working memory tasks. We examined how effectively stimulus information was represented from neurons that do and do not generate persistent activity. We also compared working memory for spatial location and object information.

## METHODS

### Animals

Data obtained from male rhesus monkeys (*Macaca mulatta*), ages 5–9 years old, weighing 5–12 kg, as previously documented (18), were analyzed in this study. None of these animals had any prior experimentation experience at the onset of our study. Monkeys were either single-housed or pair-housed in communal rooms with sensory interactions with other monkeys. All experimental procedures followed guidelines set by the U.S. Public Health Service Policy on Humane Care and Use of Laboratory Animals and the National Research Council’s Guide for the Care and Use of Laboratory Animals and were reviewed and approved by the Wake Forest University Institutional Animal Care and Use Committee.

### Experimental setup

Monkeys sat with their heads fixed in a primate chair while viewing a monitor positioned 68 cm away from their eyes with dim ambient illumination and were required to fixate on a 0.2° white square appearing in the center of the screen. During each trial, the animals maintained fixation on the square while visual stimuli were presented either at a peripheral location or over the fovea, in order to receive a liquid reward (typically fruit juice). Any break of fixation immediately terminated the trial, and no reward was given. Eye position was monitored throughout the trial using a non-invasive, infrared eye position scanning system (model RK-716; ISCAN, Burlington, MA). The system achieved a < 0.3° resolution around the center of vision. Eye position was sampled at 240 Hz, digitized and recorded. The visual stimulus display, monitoring of eye position, and synchronization of stimuli with neurophysiological data were performed with in-house software implemented on the MATLAB environment (Mathworks, Natick, MA), utilizing the Psychophysics Toolbox (19).

### Behavioral task

Monkeys were trained to complete spatial and shape working memory tasks. The stimuli shown in the spatial task were white 2° squares, presented in one of nine possible locations arranged in a 3 × 3 grid with 10° distance between adjacent stimuli. Here, we analyzed responses to only 8 stimuli, shown in the periphery of the grid (thus omitting analysis of the central, foveal stimulus). The stimuli shown in the shape task were white 2° geometric shapes, drawn from a set comprising a circle, diamond, the letter H, the hashtag symbol, the plus sign, a square, a triangle, and an inverted Y-letter. These stimuli could also be presented in one of nine possible locations arranged in a 3 × 3 grid with 10° distance between adjacent stimuli.

Presentation began with a fixation interval of 1 s where only the fixation point was displayed, followed by 500 ms of stimulus presentation (referred to hereafter as cue), followed by a 1.5 s delay interval where, again, only the fixation point was displayed. A second stimulus (referred to hereafter as sample) was subsequently shown for 500 ms. In the spatial task, this second stimulus would be either identical in location to the initial stimulus, or diametrically opposite the first stimulus. In the feature task, this second stimulus would appear in the same location to the initial stimulus and would either be an identical shape or not. Only one nonmatch stimulus was paired with each cue, so that the number of match and nonmatch trials were balanced in each set. In both the spatial and feature task, this second stimulus display was followed by a second delay period of 1.5 s where only the fixation point was displayed. The monkeys were thus required to remember the spatial location and/or shape of the first presented stimulus, and then, after the second delay period, would report whether the second stimulus was identical to the first or not, by making an eye movement to one of two target stimuli (green for matching stimuli, blue for non-matching) that appeared after the second delay period. Each target stimulus could appear at one of two locations orthogonal to the cue/sample stimuli, pseudo-randomized in each trial, so that the location of the correct target could not be predicted before the end of the trial.

A conjunction task was also used, which combined the active spatial and feature tasks, using the same shape stimuli and presented at the same possible locations, with the same timing. In a single recording session, only four shape-location combinations involving two shapes and two locations were used. The conjunction task was the most complex task in the current study, as the monkeys were required to match two different types of information: location and shape.

### Surgery and neurophysiology

A 20 mm diameter craniotomy was performed over the PFC and a recording cylinder was implanted over the site. The location of the cylinder was visualized through anatomical magnetic resonance imaging (MRI) and stereotaxic coordinates post-surgery. For two of the four monkeys that were trained to complete active spatial, feature and conjunction WM tasks, the recording cylinder was moved after an initial round of recordings in the post-training phase to sample an additional surface of the PFC.

### Anatomical localization

Each monkey underwent an MRI scan prior to neurophysiological recordings. Electrode penetrations were mapped onto the cortical surface. We recorded from a posterior-dorsal region that included area 8A, a mid-dorsal region that included area 8B and area 9/ 46, an anterior-dorsal region that included area 9 and area 46, a posterior-ventral region that included area 45, and an anterior-ventral region that included area 47/12.

### Neuronal recordings

Neural recordings were carried out in the aforementioned areas of the PFC both before and after training in each WM task. Subsets of the data presented here were previously used to determine the collective properties of neurons in the dorsal and ventral PFC, as well as the properties of neurons before and after training in the posterior-dorsal, mid-dorsal, anterior-dorsal, posterior-ventral, and anterior-ventral PFC subdivisions. Extracellular recordings were performed with multiple microelectrodes that were either glass- or epoxylite-coated tungsten, with a 250 μm diameter and 1–4 MΩ impedance at 1 kHz (Alpha-Omega Engineering, Nazareth, Israel). A Microdrive system (EPS drive, Alpha-Omega Engineering) advanced arrays of up to 8-microelectrodes, spaced 0.2–1.5 mm apart, through the dura and into the PFC. The signal from each electrode was amplified and band-pass filtered between 500 Hz and 8 kHz while being recorded with a modular data acquisition system (APM system, FHC, Bowdoin, ME).

Waveforms that exceeded a user-defined threshold were sampled at 25 μs resolution, digitized, and stored for off-line analysis. Neurons were sampled in an unbiased fashion, collecting data from all units isolated by our electrodes, with no regard to the response properties of the isolated neurons. A semi-automated cluster analysis relied on the KlustaKwik algorithm, which applied principal component analysis of the waveforms to sort recorded spike waveforms into separate units. To ensure a stable firing rate in the analyzed recordings, we identified recordings in which a significant effect of trial sequence was evident at the baseline firing rate (ANOVA, p < 0.05), e.g., due to a neuron disappearing or appearing during a run, as we were collecting data from multiple electrodes. Data from these sessions were truncated so that analysis was only performed on a range of trials with stable firing rate. Less than 10% of neurons were corrected in this way. Identical data collection procedures, recording equipment, and spike sorting algorithms were used before and after training in order to prevent any analytical confounds.

### Data analysis

Data analysis was implemented with the MATLAB computational environment (Mathworks, Natick, MA). Neuron analysis involved, initially, determining the mean firing rate of each neuron in each trial epoch. Next, we compared responses of each neuron in the 1 s fixation period with each of the task epochs that followed (cue presentation, first delay period, sample presentation, second delay period). Any neuron that displayed a significantly greater response in a particular epoch over the fixation was identified as a task-responsive neuron (one-tailed paired t-test; p < 0.05). We particularly focused on the neurons with significantly elevated delay period activity in the first delay period. These are the neurons classically defined as having persistent activity (20, 21).

Decoding analysis was performed as follows: Spiking responses from 1 s before cue onset to 5 s after cue onset were binned using a 400-ms-wide window and 100 ms steps to create a spike count vector with 57 elements. We additionally examined decoder performance for a 1 s interval in the center of the delay period (250 ms after the cue offset). Spike count vectors from multiple neurons were used to create pseudo-populations of 50, 100, and 200 randomly selected neurons with persistent activity or not. Five decoding algorithms were implemented using the MATLAB fitc functions to decode stimuli location: support vector machine (SVM) with a gaussian kernel, SVM with a linear kernel, discriminant analysis, classification trees, and naive bayes. Decoder performance was estimated via 10-fold and 5-fold cross-validation methods. The decoding baseline was 12.5% for both tasks as we used 8 possible locations for the spatial task and 8 possible shapes for the feature task.

## RESULTS

Extracellular neurophysiological recordings were collected from the lateral PFC of three monkeys trained to perform the match/nonmatch WM tasks (18, 22). The task required them to view two stimuli appearing in sequence with an intervening delay period between them and, after a second delay period, to report whether or not the second stimulus was identical to the first. The two stimuli could differ in terms of their location (spatial task, Fig. 1A), shape (feature task, Fig. 1B), or both (conjunction task, Fig. 1C). If the second stimulus matched with the first, monkeys would saccade towards a green target during a subsequent interval. Otherwise, they would saccade to a blue target at the diametrically opposite location.

**Figure 1.**
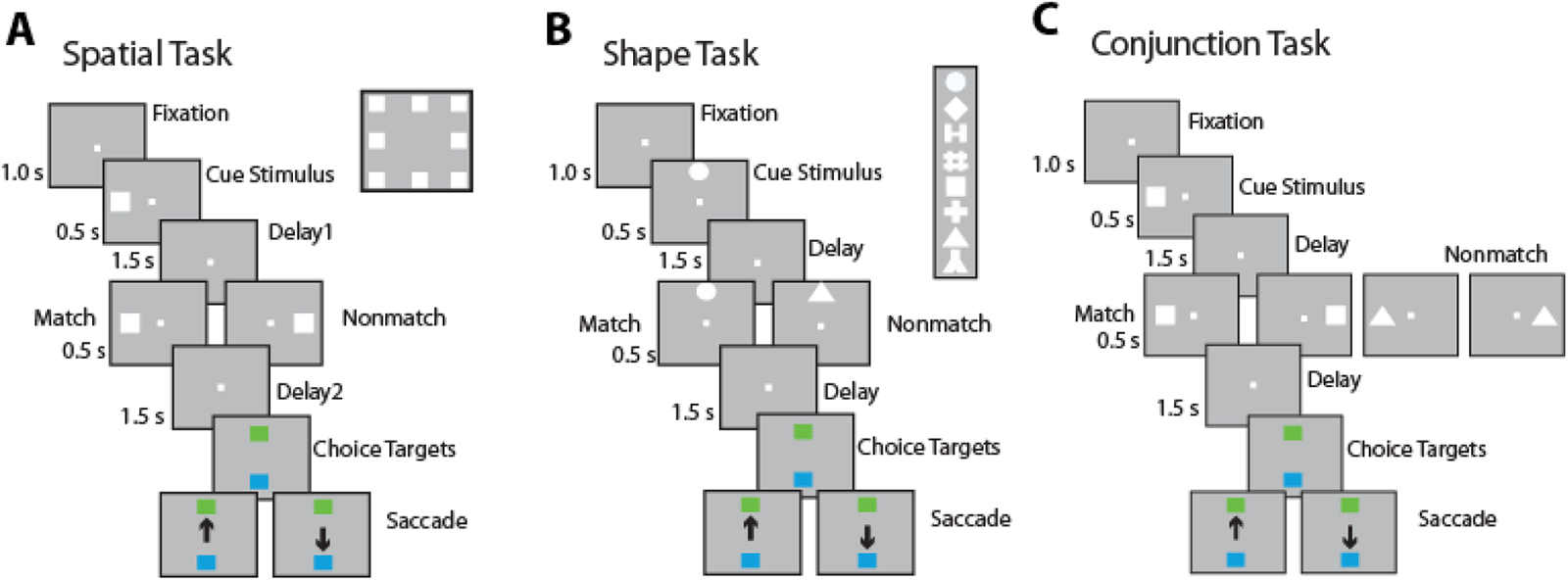
Task structure and stimuli used. The animals were required to maintain center fixation throughout the trial. When the fixation point was turned off, monkeys were required to make a saccade to a green target if two the stimuli presented were identical (matched) or to a blue target if the stimuli were different (did not match). (A) Spatial location match-nonmatch task. Eight cue locations used for analysis are shown in the inset. (B) Shape feature match-to-sample task. Eight shapes used for analysis are shown in the inset. (C) Spatial-shape conjunction task. Two locations and two stimulus shapes were used for any single session.

A total of 1066 cells from three monkeys and 885 cells from two monkeys were collected while the animals were performing the spatial and shape tasks, respectively and had sufficient numbers of trials for analysis. In the spatial task, 414 neurons exhibited persistent activity in the delay period (“persistent neurons” for simplicity, thereafter). We additionally identified 207 task-responsive, but non-persistent neurons (i.e. neurons that had significantly activity in a stimulus presentation period, but not the delay period). Another 371 neurons exhibited no significantly elevated activity in any task interval. We refer to the latter two groups of 578 neurons as “all non-persistent” neurons). Finally, a group of 74 neurons exhibited significantly inhibited activity during the delay period of the task, as we have described previously (23); these were analyzed separately.

The grant average firing rate across all neurons and stimulus conditions in the spatial task revealed only a modest increase in firing during the stimulus presentation periods (which are represented by the black horizontal lines at the bottom of Fig. 2A). Elevated activity in the delay period of the task is barely noticeable in this plot, as most prefrontal neurons do not generate persistent activity. Delay period activity is more noticeable in the neurons that generate persistent activity (red trace in Fig. 2A), but even for those it is important to note that persistent activity is typically tuned, so that each neuron generates persistent activity for only a narrow range of locations (20) and averaging responses across all stimulus conditions will inevitably mask this stimulus-selective activity (21).

**Figure 2.**
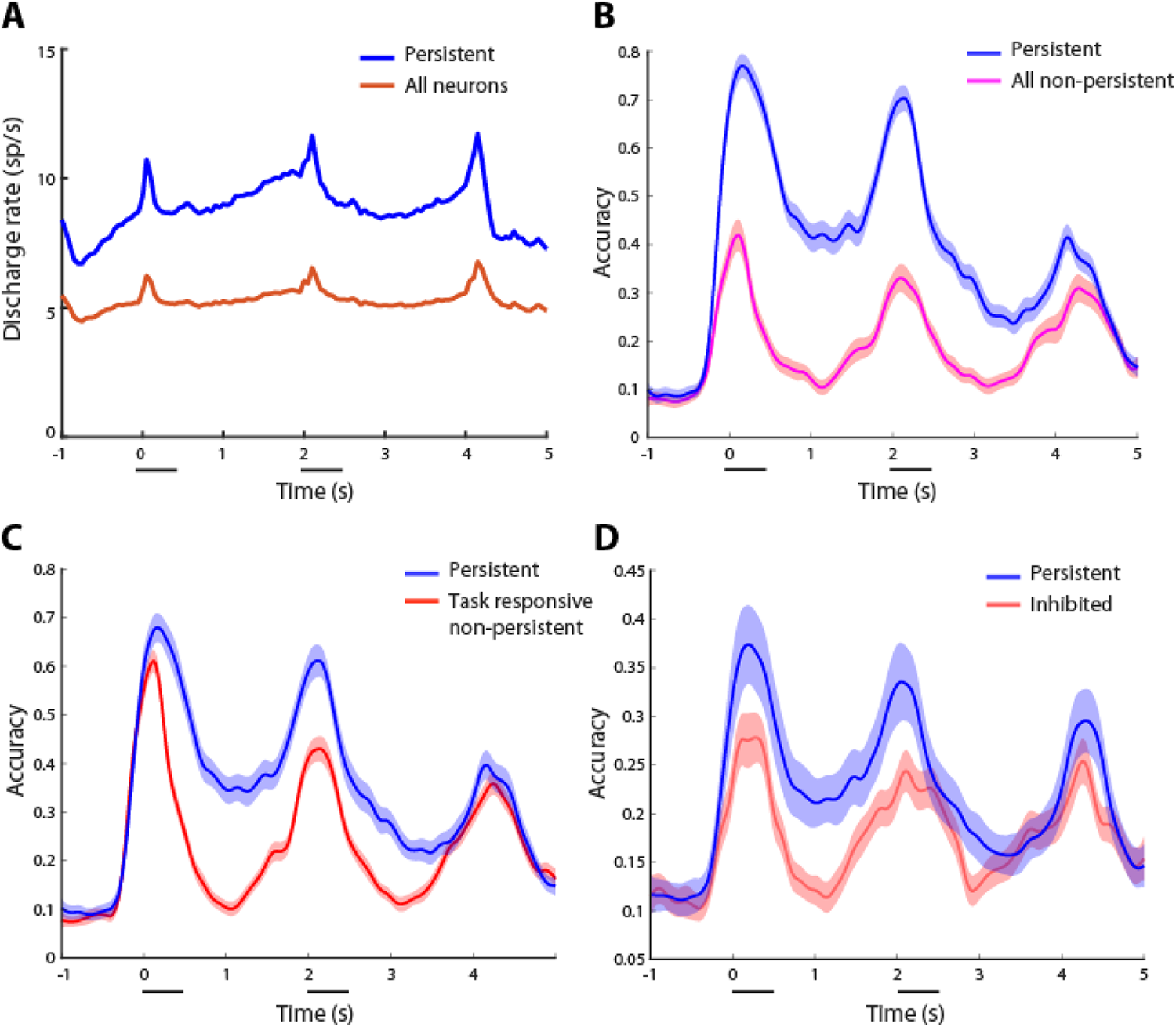
(A) Average, population Peri-Stimulus Time Histogram (PSTH) for all recorded neurons, and for neurons that generated persistent activity (persistent neurons) in the spatial working memory task. (B) SVM decoding performance for the spatial task as a function of time relative to the onset of the cue. Accuracy for populations of persistent and all non-persistent neurons with n=200 neurons and 10-fold cross-validation. Shaded area represent standard deviation calculated over 100 repetitions. (C) Decoding accuracy for populations of persistent and task-responsive but non-persistent neurons (n=200 neurons). (D) Decoding accuracy for populations of persistent and inhibited neurons (n=50 neurons).

### Decoding information from persistent and non-persistent neurons

We compared the classification accuracy of Support Vector Machine (SVM) decoding for task responsive neurons with and without persistent delay period activity across the time course of the spatial match-non match task (Fig. 2B). Mean classifier performance for decoding the location of the first (cue) stimulus with 10-fold cross-validation is shown in Fig. 2B. Persistent neurons exhibited higher classification performance at all task intervals, including the delay period. By contrast, classification accuracy essentially returned to chance levels at about 0.125 s after the beginning of the delay period for non-persistent neurons.

We also compared the classification accuracy of persistent neurons against the accuracy of task-responsive but non-persistent neurons, using equal sample sizes (Fig. 2C). Both populations reached a similar peak shortly after the presentation of the first (cue) stimulus, indicated by the horizontal line in Fig. 2. However, decoding accuracy diverged during the delay period, with the largest difference between populations being seen during the first delay period (0.5-2.0s). A similar difference in decoding performance was also present during the second delay period, although it should be noted that at this point in the task, maintenance of the first stimulus appearance is no longer required; the monkeys could make a judgement about whether or not the two stimuli were matching after the appearance of the second stimulus, without needing to continue maintaining the location of either the first or second stimulus in memory. Neurons with significantly reduced delay period activity also failed to match the decoding performance of persistent neurons (inhibited neurons in Fig. 2D). The analysis that follows involves primarily comparisons of persistent neurons with task-responsive, non-persistent neurons, to better understand the difference in properties in the delay period for neurons that carry comparable information during the stimulus presentation period.

To determine whether persistent and task-responsive, non-persistent neurons could maintain spatial location information with equal accuracy during the delay period, and to avoid neurons with potentially long response latencies that might carry over at the first part of the delay interval, we calculated SVM decoding accuracy for the middle of the first delay period (0.75-1.75s). Mean decoding accuracy was higher for persistent populations (0.47) than non-persistent populations (0.17) for 100 trials (Fig. 3), with essentially no overlap between the two distributions. A statistical comparison based on 100 simulations for each condition revealed a significant difference (t-test, *t_198_*=72.6, p<0.001). This suggests that persistent neurons encode information substantially more accurately during the delay period. However, it was also notable that non-persistent neurons did achieve a classification accuracy greater than chance in this period (t-test, *t_99_*=21.5, p<0.001).

**Figure 3.**
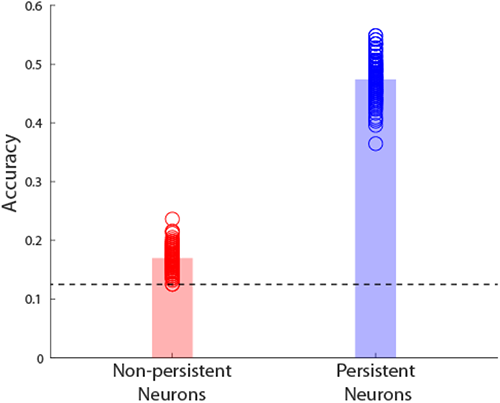
SVM decoding performance for the spatial task. Accuracy for the middle 1 s of the first delay period. Simulations are shown separately for persistent and (task-responsive) non-persistent neurons with n=200 neurons and 10-fold cross-validation. Horizontal line represents chance performance.

### Effect of sample size and classifier parameters

The absolute performance achieved by classifiers depends on the amount of neural data used, as well as the choice of parameters. It was therefore important to examine whether the difference between persistent and non-persistent neurons was sensitive to these factors. We first varied the population size. Populations of 50, 100, and 200 neurons showed time courses consistent with the spatial task with the SVM decoding algorithm. Higher decoding accuracy for persistent populations during the first delay period was consistent across population sizes (Fig. 4A). We saw increased accuracy at the first timepoint in the delay period as a function of the number of neurons, for both persistent and non-persistent populations (Fig. 4B). However, persistent neurons achieved consistently higher performance.

**Figure 4.**
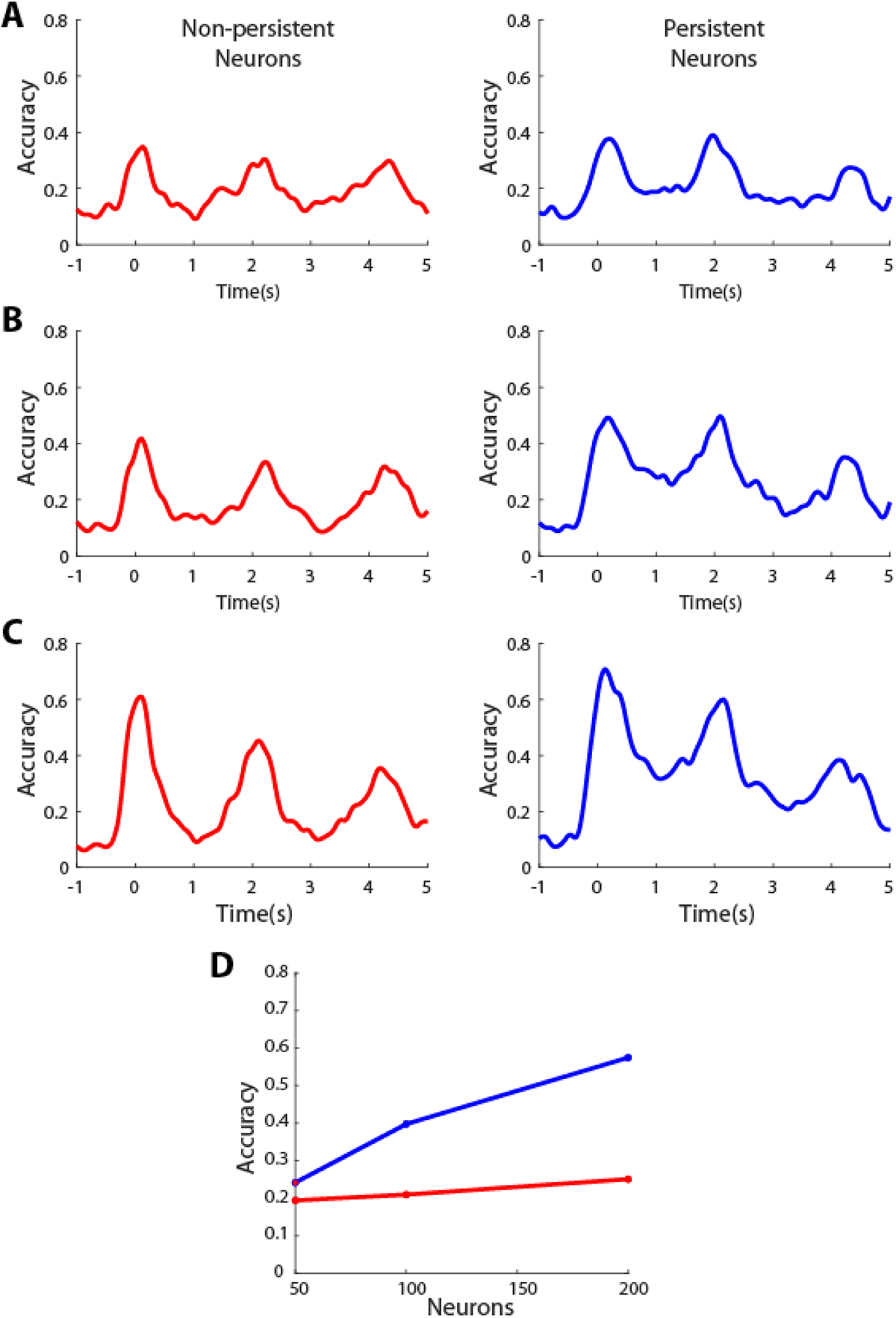
SVM decoding performance in the spatial task, for different sample sizes of persistent and (task-responsive) non-persistent neurons. (A-C). Sample size of n=50, 100, and 200 neurons respectively. (D). Accuracy of data shown for the first time point of the delay period.

Decoding performance over time for the spatial task with the SVM decoder was also consistent between 10-fold and 5-fold cross-validation (Fig. 5). For both non-persistent and persistent populations, we once again saw the maxima when the stimuli were present, and saw the minima during the delay periods when the stimuli were not present. Higher accuracy during delay periods for persistent neurons was seen for both 10-fold and 5-fold cross-validation.

**Figure 5.**
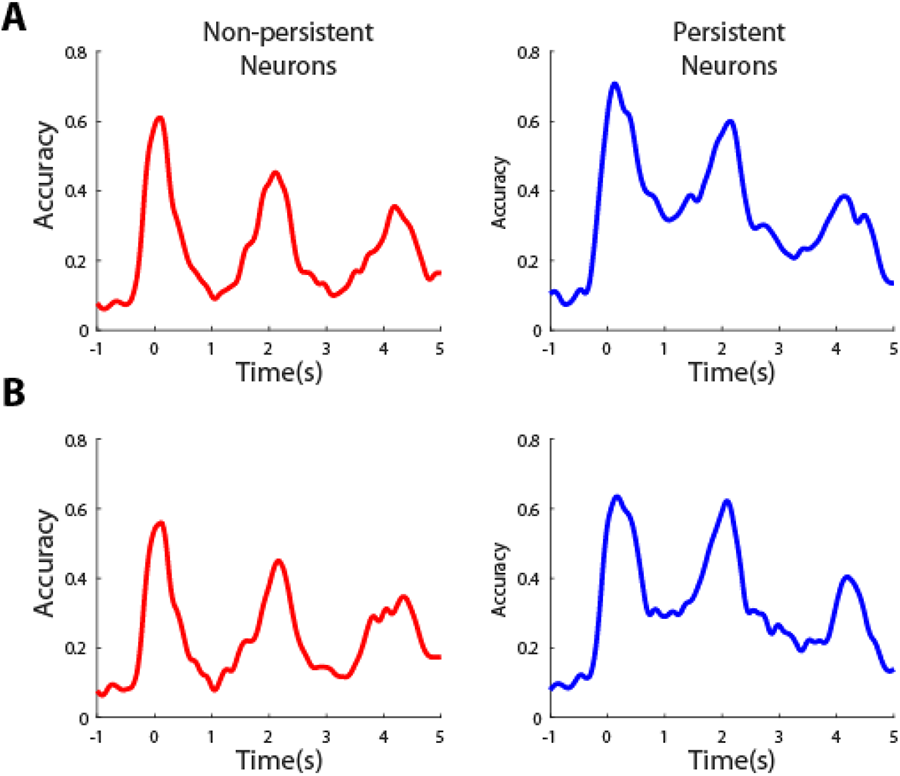
SVM decoding performance in the spatial task, for different parameters of cross-validation for persistent and (task responsive) non-persistent neurons. (A) Five-fold cross validation. (B). Ten-fold cross validation. Sample size n=200 neurons, for all cases.

Finally, we compared the performance of different classifiers against each other, including SVM, Discriminant Analysis, Naïve Bayes, and Classification trees (Fig. 6). SVM generally performed the best in our dataset. However, for every classifier, persistent neurons consistently outperformed non-persistent neurons in the delay period of the task.

**Figure 6.**
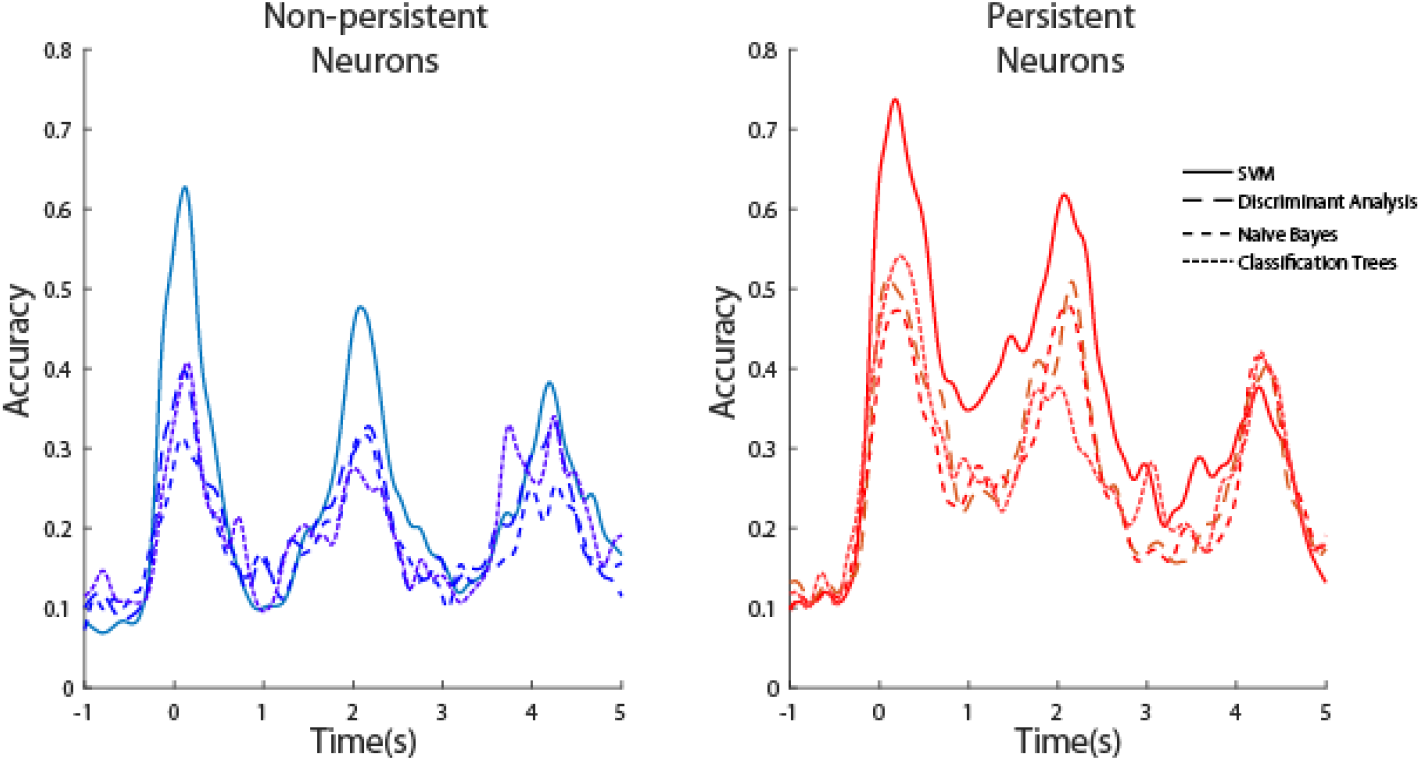
Decoding performance in the spatial task, for persistent and (task responsive) non-persistent neurons, using different classifiers. Sample size n=200 neurons, for all cases.

### Decoding object working memory

The results of the prior sections suggest that neurons that generate persistent activity can maintain spatial working memory better than those without persistent activity. However, alternative theories of working memory have argued that the role of the prefrontal cortex is to “highlight” the spatial location of remembered stimuli, perhaps playing a supervisory or control role, while sensory areas maintain the actual features of stimuli in memory (24–27). Indeed, the findings that put forth activity-silent theories relied on the general lack of increased persistent activity during the delay periods of object working memory tasks (10, 11). It was therefore critical to compare results from our shape working memory task, which required monkeys to remember the shapes of stimuli, that were always presented at the same location (Fig. 1B).

Similarly to the spatial task, we identified 315 neurons that generated persistent activity (“persistent neurons”), 183 neurons that responded to the task but did not generate persistent activity, and 312 neurons that did not respond to the task. The latter two groups again made up 495 “all non-persistent” neurons. Finally, we identified 75 neurons whose activity was inhibited in the delay period. As was the case for the spatial task, the grant average firing rate across all neurons and stimulus conditions revealed very little delay period activity (Fig. 7A). This result essentially replicates the findings of previous studies that reported lack of increased persistent activity during the delay periods of object working memory tasks (10, 11). As we explained above, this apparent lack of delay activity is due to the fact that only a proportion of cells show persistent activity in the whole population (35.6% in our shape working memory task). Additionally, those cells are generally selective and only increase their firing rates to one or few particular stimuli rather than to all stimuli used. Thus, a grant average will mask the elevated activity of persistent cells.

**Figure 7.**
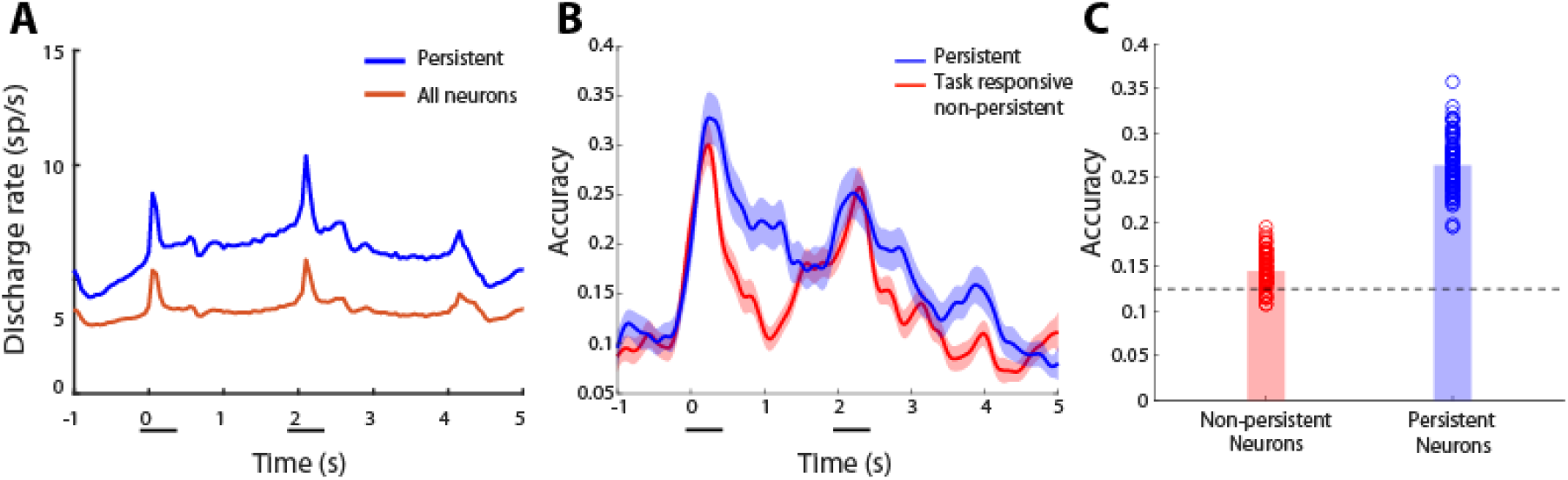
(A) Average PSTH for all recorded neurons and persistent neurons in the shape working memory task. (B) Decoding performance in the shape working memory task, for persistent and (task responsive) non-persistent neurons, using the SVM classifier, with a 10-fold cross-validation. Sample size n=180 neurons, for both cases. Time course of decoding accuracy during the duration of the trial. Conventions are the same as in Fig. 2. (C) Decoding accuracy for (task responsive) non-persistent neuros during the middle 1 s of the delay period (0.75 – 1.75 s). Conventions are the same as in Fig. 3.

Results of SVM decoder classification accuracy are shown in Fig. 7B. Persistent and non-persistent neurons reached similar classification accuracy during the stimulus presentation period. Importantly, persistent neurons consistently outperformed non-persistent neurons during the delay periods of the task (Fig. 7C). A statistical comparison based on 100 simulations for each condition revealed a significant difference (t-test, *t_198_*=33.1, p<0.001). This suggests that persistent neurons encode object information substantially more accurately during the delay period. In this case, too non-persistent neurons did achieve a classification accuracy greater than chance in this period (t-test, *t_99_*=10.6, p<0.001). In this respect, the results of the spatial and object working memory tasks were qualitatively similar, although we note that the absolute value of classification was lower in the object task compared to the spatial task, as a result of fewer neurons being engaged by any shape stimulus.

### Decoding both spatial and shape information from working memory

We have so far investigated the cases where one type of information was required to be maintained in the working memory, and revealed that the persistent neurons contain more information than the nonpersistent cells. However, it is still unknown if this is true in more complex situations, like when combinations of different features are remembered. We thus analyzed 273 persistent neurons and 141 nonpersistent (task responsive) neurons in the conjunction task (Fig. 1C). Results are shown in Fig. 8. Again, the result is qualitatively similar compared to the scenario when only a single type of information was remembered. Despite of similar level of decoding during the stimulation presentation epochs (the cue and sample period), cells without persistent activity in the delay period contain significantly less information of stimuli identity during the delay period (t-test, *t_198_*=41.9, p<0.001).

**Figure 8.**
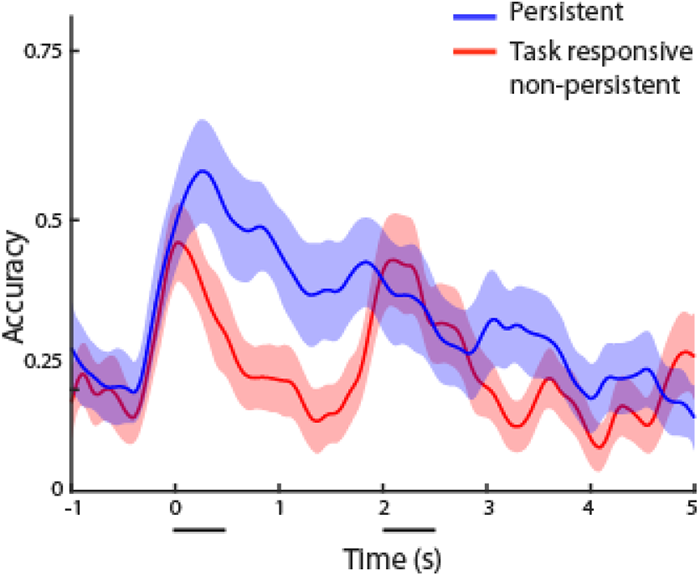
Decoding performance in the conjunction working memory task, for persistent and (task responsive) nonpersistent neurons, using the SVM classifier, with a 10-fold cross-validation. The stimuli identity (cue-shape combination) in the cue period was decoded. Sample size n=100 neurons, for both cases. Time course of decoding accuracy during the duration of the trial. Conventions are the same as in Fig. 2B.

## DISCUSSION

The role of persistent activity has been reevaluated recently (10, 28, 29). Although strong evidence links persistent activity of prefrontal neurons with behavior in some working memory tasks (21), objections have been raised about a general role of persistent activity across all working memory functions. It has thus been suggested that persistent activity in spatial tasks may represent motor preparation rather than the remembered spatial location of the stimuli per se (9). Alternatively, the prefrontal cortex may play a supervisory role, “highlighting” the locations of stimuli, but not maintaining information about object identity (26, 30, 31).

Our study identified neurons that generate readily persistent activity in a spatial working memory task that separated the stimulus location from the eventual motor response. Our results showed that most information about the remembered stimulus location—which was necessary to perform the match-nonmatch task—was maintained by neurons that generated persistent activity. This persistent activity was also shown to be informative about the identity of stimuli held in memory, not just their spatial location. Our results therefore demonstrate that persistent activity is the primary mechanism of information maintenance across working memory stimulus domains in the primate prefrontal cortex.

### Information decoding from persistent and non-persistent neurons

Our analysis identified neurons with persistent activity, classically defined as activity that is elevated by a statistical criterion in the delay period compared to the fixation period of the task (20). The visual displays that our subjects observed during these two task epochs were identical, with only the fixation point appearing on-screen (see Fig. 1). The difference between these epochs thus lay in the stimulus presentation between them, which the subjects were taught to maintain in memory. Neurons that generated persistent activity accounted for most of the spatial and object information that could be decoded by the prefrontal population during the delay period. The effect was consistent across differently sized sets of neurons used to implement classification, across different classification parameters and across different kinds of classifiers. An important caveat for this analysis is that neurons without persistent activity still exhibited some information, as predicted by activity silent theories (10). On the other hand, it is also possible that some of the neurons which failed to meet our statistical criterion, generated some traces of persistent activity.

Neurons with persistent activity comprised the minority of neurons in the shape working memory task. In fact, overall neuronal activity, averaged across all neurons and stimuli, barely suggested increased firing rate in the delay period of the task. However, classification based on non-persistent neurons barely exceeded chance performance. This result thus provides direct evidence in favor of the idea that prefrontal neurons maintain stimulus related information in working memory by virtue of persistent discharges (32).

### Activity Silent Mechanisms

Strong theoretical and experimental evidence exists for some types of activity-silent mechanisms operating during working memory tasks. For example, activity elicited after repeated presentation of the same stimulus is typically reduced for many neurons, a phenomenon termed repetition suppression, even though persistent activity may not be present in most of these neurons (33). The response difference to matching and nonmatching stimuli has predictive power over behavioral performance, differing systematically in correct and error trials, and thus providing evidence of memory being influenced by this activity (7, 34). Computational models have been proposed that could account for such changes via mechanisms that depend on synaptic strength modifications instead of spike generation (12, 13). Recurrent Neural Networks relying on rapid changes in synaptic weights after a stimulus presentation are able to perform working memory tasks (14–17). However, these studies have also suggested that synaptic mechanisms alone are inadequate to maintain information for more demanding tasks, such as tasks involving multiple sequential stimuli (15).

Serial bias is another phenomenon attributed to silent mechanisms. Specifically, the location of a previous trial’s stimulus biases the response in the current trial. Recent spatial working memory studies have suggested this mechanism to be both present and compatible with persistent activity mechanisms. When the appearance of a fixation point increases neural excitability prior to the stimulus presentation, the previous trial’s activity bump could undergo a transient reactivation, which may then merge with the new bump (35). In essence, synaptic mechanisms influence working memory to the extent that they influence spiking generation (32).

### Working memory for complex tasks

Prefrontal neurons generate persistent discharges in memory tasks that require the maintenance of stimulus features such as shape, color, direction of motion (36–41), the identity of objects, faces, and abstract pictures (42–49). These findings are also complicated by the fact that prefrontal persistent activity represents information beyond the characteristics of stimuli, including the rules of the cognitive task (50, 51), categories (44, 52), numerical quantities (53) and other abstract factors. It has thus been suggested that persistent activity may be mostly driven by task processes, rather than stimulus maintenance (54). Indeed, training to perform working memory tasks generally increases firing rate, particularly in delay periods, and induces a variety of other changes in factors such as variability and correlation (55, 56).

Our current results argue against the interpretation that the activity of prefrontal neurons is entirely representing task and rule information. On the contrary, we identified stimulus selective, persistent activity that maintained information about the remembered stimuli themselves during the delay period of the task within the prefrontal cortex. Generation of persistent activity in the prefrontal cortex thus appears to be a critical mechanism of working memory maintenance.

## ACKNOWLEDGMENTS

Research reported in this paper was supported by the NIH National Eye Institute under award numbers R01 EY017077 and P30 EY008126 and by the National Institute of Mental Health under award number R01 MH116675. We wish to acknowledge Chrissy Suell for technical help; Kai Malcolm for initial simulations that led to this project; and Junda Zhu, and Zhengyang Wang for helpful comments on the manuscript.

